# Enhancing NeuroD1 Expression to Convert Lineage-Traced Astrocytes into Neurons

**DOI:** 10.1101/2022.06.21.496971

**Authors:** Liang Xu, Zong-Qin Xiang, Yao-Wei Guo, Yu-Ge Xu, Min-Hui Liu, Wen-Yu Ji, Shu He, Wen-Liang Lei, Wen Li, Zheng Wu, Gong Chen

## Abstract

Regenerating functional new neurons in adult mammalian brains has been proven a difficult task for decades. Recent advancement in direct glia-to-neuron conversion *in vivo* opens a new field for neural regeneration and repair. However, this emerging new field is facing serious challenges from misuse of viral vectors to misinterpretation of conversion data. Here, we employ a variety of AAV vectors with different promoters and enhancers to demonstrate that astrocytes can be converted into neurons in a NeuroD1 dose-dependent manner in both wildtype (WT) and transgenic mice. Notably, astrocytes in WT mice were relatively easy to convert with higher conversion efficiency, whereas lineage-traced astrocytes in Aldh1l1-CreERT2 mice showed high resistance to reprogramming but were still converted into neurons after enhancing NeuroD1 expression with CMV enhancer. Furthermore, under two-photon microscope, we observed direct astrocyte-to-neuron conversion within 3 weeks of serial live imaging in the mouse cortex. We also demonstrated that high titre AAV reaching 10^13^ GC/ml caused severe neuronal leakage using a variety of AAV GFAP::GFP vectors, highlighting the necessity to inject low titre AAV into healthy brains to avoid artifactual results. Together, our studies suggest that lineage-traced astrocytes can be converted into neurons but require stronger conversion force such as enhanced NeuroD1 expression. Failure to recognize the difference between WT astrocytes and lineage-traced astrocytes in terms of conversion barrier will lead to misinterpretation of data.

## INTRODUCTION

Brain repair in adult mammalian animals is very challenging due to lack of neurogenesis after injury or diseases. Transplantation of externally cultured cells into the central nervous system (CNS) may produce certain number of new neurons to treat specific neurological disorders such as Parkinson’s disease (Schweitzer et al., 2020; Tao et al., 2021). An alternative approach for regenerating new neurons is to convert internal glial cells into functional neurons (Barker et al., 2018; Bocchi et al., 2021; Lei et al., 2019; Li and Chen, 2016; Qian and Fu, 2021). Such direct in vivo reprogramming technology has been demonstrated repeatedly in my lab (Chen et al., 2020; Ge et al., 2020; Guo et al., 2014; Liu et al., 2020; Puls et al., 2020; Wu et al., 2020; Xiang et al., 2021; Zhang et al., 2020; Zheng et al., 2022), as well as in many other labs around the world (Gascon et al., 2016; Heinrich et al., 2014; Jiang et al., 2021; Lentini et al., 2021; Liu et al., 2015; Niu et al., 2015; Niu et al., 2013; Qian et al., 2020; Su et al., 2014; Tang et al., 2021; Torper et al., 2015). In vivo reprogramming is not only achieved in the CNS and but also in other organ systems such as pancreas, heart, and liver (Larouche and Aguilar, 2019; Nam et al., 2014; Qian et al., 2012; Rezvani et al., 2016; Song et al., 2016; Zhou et al., 2008). However, a few studies recently raised concerns regarding direct glia-to-neuron (GtN) conversion in the CNS, reporting lack of glial conversion in astrocytic lineage-tracing Aldh1l1-CreERT2 mice (Hoang et al., 2021; Wang et al., 2021). Without understanding the conversion barrier in WT astrocytes versus lineage-traced astrocytes, some researchers quickly jumped on a conclusion that previous conversion studies might be due to reporter leakage into pre-existing neurons. These assertions had caused huge confusion in the conversion field (Chen, 2021). Are previous studies published by so many different labs all wrong?

To find the answer, we designed a series of experiments to address critical points that have been misleading the glial conversion field over the past year. The first mis-concept and a common mistake in glial conversion field is that one can freely inject high titre viral vectors into normal healthy brains without worrying about adversary effects. Here, we demonstrated that once reaching high titre of 1×10^13^ GC/ml at 1 μl in the mouse cortex, control AAV vectors (GFAP::GFP) started to express GFP directly in neurons, leading to severe neuronal leakage. The second mis-concept about glial conversion is that only lineage-traced glial cells can be adopted as the golden standard for glial conversion. We demonstrated that compared to WT astrocytes, lineage-traced astrocytes were much more difficult to convert. However, when we used CMV enhancer to increase NeuroD1 expression in the lineage-traced astrocytes of Aldh1l1-CreERT2 mice, we detected clear neuronal conversion from lineage-traced astrocytes. Therefore, lineage-tracing mice cannot be used as golden standard for glial conversion, because they have much higher conversion barrier to overcome. Understanding the high conversion barrier in the lineage-traced astrocytes is critical for the entire glial conversion field, because it demands much greater conversion force such as enhancing the expression level of transcription factor(s) (TF) to overcome the conversion barrier. It is not a surprise that those who are only using weak GFAP promoter to drive TF expression have failed to convert lineage-traced astrocytes into neurons. Once they enhance the conversion force to overcome the conversion barrier, the lineage-traced astrocytes will be converted into neurons.

## RESULTS

### High titre AAV causing severe neuronal leakage

For in vivo astrocyte-to-neuron conversion, many labs have now used AAV instead of retrovirus because of the high transduction efficiency of AAV in the CNS. However, different labs are using different AAV titres, ranging widely from 10^8^ – 10^14^ GC/ml, in conducting glial conversion studies and hence reaching different conclusions. To solve this chaotic issue regarding AAV titre (volume typically 1-2 μl, not different among different labs), we tested 3 different titres (10^11^, 10^12^, 10^13^ GC/ml; volume kept the same at 1 μl) using 3 AAV vectors (GFAP2.2::GFP, GFAP1.6::GFP, and GFAP104::GFP) that express GFP typically in astrocytes (Fig. 1A). At 30 days post viral injection (dpi), brain samples were collected and immunostained to analyze GFP expression in astrocytes and neurons (Fig. 1B). We found that all 3 vectors expressed GFP efficiently in the mouse cortex at all 3 titres, and GFP expression showed dose-dependent increase as expected (Fig. 1C-E). It is interesting to note that while all 3 vectors infected cortical columns from superficial layer I to deep layer VI at the injection site, we did observe much wider AAV spreading in the superficial layer I-III (Fig. 1C-E). We speculate that such AAV spreading pattern might be related to the blood vessel and lymphatic tube trajectories in the superficial layer of cortex (Murlidharan et al., 2016). Immunostaining with astrocytic marker S100b revealed that low titre AAV expressed GFP essentially all in astrocytes (Fig. 1F-H, 10^11^ GC/ml columns, white arrowhead), but at high titre of 10^13^ GC/ml, many GFP-expressed cells did not show S100b signal but displayed neuronal morphology (Fig. 1F-H, 10^13^ GC/ml columns, yellow arrowhead). Immunostaining with neuronal marker NeuN revealed that low titre AAV (10^11^ GC/ml) did not express GFP in NeuN+ neurons, but high titre AAV (10^13^ GC/ml) directly expressed GFP in NeuN+ neurons although GFP expression was controlled by astrocytic promoter GFAP (Fig. 1I-K, red arrowhead), suggesting a significant leakage of high titre AAV into neurons. Quantitative analyses showed that low titre AAV at 10^11^ GC/ml did not leak into neurons regardless of the GFAP promoters (Fig. 1L-N, gray bars), but high titre AAV at 10^13^ GC/ml resulted in severe neuronal leakage for all 3 GFAP promoters (Fig. 1I-K, blue bars, leakage rate 50-60%). The moderate titre of 10^12^ GC/ml showed moderate GFP leakage into neurons with large variations among different promoters (Fig. 1L-N, green bars). Therefore, regardless of the restriction of GFAP promoter, high titre AAV at 10^13^ GC/ml or above will cause severe neuronal leakage, making any data interpretation inaccurate. Based on our dose-finding studies, we recommend that glial conversion studies should use a moderate titre ranging from 10^11^ - 10^12^ GC/ml (1-2 μl for mouse brains) to avoid severe neuronal leakage and mis-interpretation of the data. Previous results using high titre AAV at ≥10^13^ GC/ml published by Wang et al. must be repeated with lower titre AAV at 10^11^ - 10^12^ GC/ml in order to correct wrong interpretations of their data (Wang et al., 2021).

**Figure 1.**
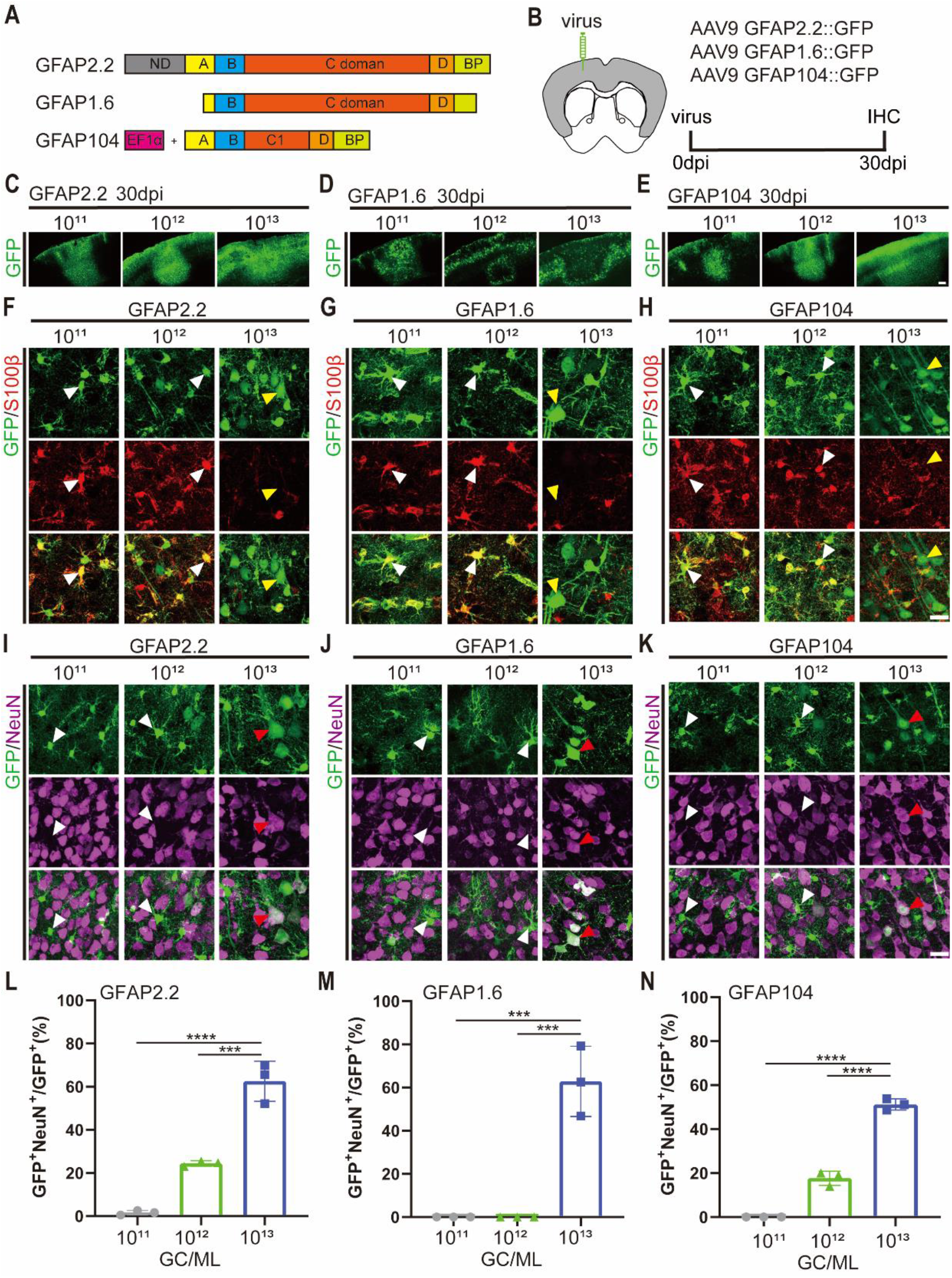
High titre AAV causes dose-dependent neuronal leakage. (A) Schematic outlines of the three selected GFAP promoters. (B) Schematic diagram showing virus injection site and experimental design. (C,D,E) Representative images showing GFP expression throughout the mouse cortices infected with AAVs with different promoters and at different titres. (F,G,H) High-magnification images showing the relationship between GFP-expressing cells (green) by different GFAP::GFP viral vectors and S100β-labeled astrocytes (red) at different AAV titres. Scale bar = 20 μm. (I,J,K) High-magnification images showing the relationship between GFP-expressing cells (green) and NeuN+ neurons (violet). Note that at high AAV titre of 10^13^ GC/ml, many GFAP::GFP-infected cells were NeuN+ neurons, indicating severe neuronal leakage. Scale bar = 20 μm. (L,M,N) Bar graphs showing the neuronal leakage rate calculated by GFP+NeuN+ cells among total GFP+ cells. The neuronal leakage rate increased significantly in an AAV titre-dependent manner. Values are shown as mean ± SD. N = 3 per group. One-way ANOVA analysis with Tukey post hoc test, ***p < 0.001, ****p < 0.0001.

### NeuroD1 dose-dependently converts astrocytes into neurons

After solving the neuronal leakage issue induced by high titre AAV, we revisited NeuroD1-mediated astrocyte-to-neuron (AtN) conversion. We used low titre (10^11^ GC/ml) AAV GFAP104::GFP to label astrocytes specifically (Fig. 1N), and then added different doses of AAV GFAP104::NeuroD1 to test their conversion capability (Fig. 2A). If NeuroD1 does not convert astrocytes at all, then we expect that GFAP104::GFP-labeled astrocytes shall remain astrocytes without any changes regardless of any NeuroD1 added. On the contrary, what we discovered was a striking reduction of GFP expression that was inversely correlated with an increase of NeuroD1 expression (Fig. 2B). Clearly, NeuroD1 expression downregulated GFP expression in GFAP104::GFP-labeled astrocytes. To understand why astrocytes lose their GFP signal in the presence of NeuroD1, we performed immunostaining of astrocytic marker GFAP and neuronal marker NeuN. As expected, control AAV GFAP104::GFP-infected cells were GFAP+ astrocytes (Fig. 2C, top row). Low titre AAV (10^11^ GC/ml) GFAP104::NeuroD1 expressed very low level of NeuroD1, and the GFP-labeled cells showed typical astrocytic morphology and colocalized with GFAP (Fig. 2C, 2^nd^ row). In contrast, after adding higher titre AAV (5×10^11^ - 5×10^12^ GC/ml) GFAP104::NeuroD1, many GFP-labeled cells lost both GFP and GFAP signal but acquired NeuN signal (Fig. 2C, 3^rd^ row and 4^th^ row, red arrowhead). Quantitative analyses showed that GFP intensity decreased from 8000 a.u. in the control group (GFAP104::GFP alone) to 4000 a.u. after adding very low titre AAV GFAP104::NeuroD1 (10^11^ GC/ml), and further decreased to <1000 a.u. after adding high titre AAV GFAP104::NeuroD1 (5×10^12^ GC/ml) (Fig. 2D). Such sharp decrease of GFP intensity was inversely correlated with an increase of NeuroD1 intensity (Fig. 2E) as well as an increase in the number of GFP+ neurons (Fig. 2F). Note that because GFP signal was greatly downregulated by NeuroD1, many astrocyte-converted neurons may not show any GFP signal any more. Thus, the conversion rate using GFP as the marker could be an underestimate of the real conversion rate. Nevertheless, these results suggest that NeuroD1 dose-dependently converts astrocytes into neurons.

**Figure 2.**
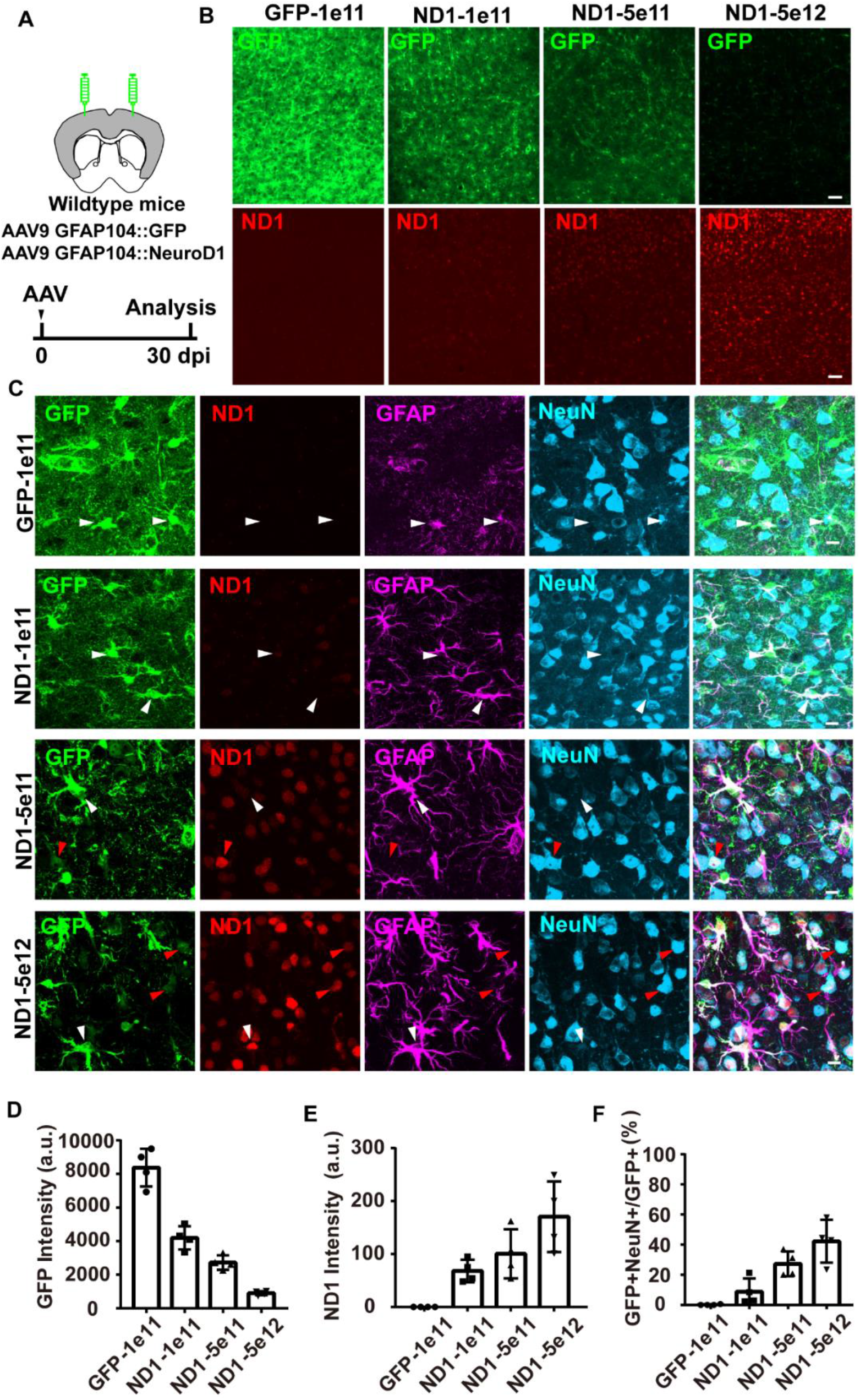
NeuroD1 dose-dependent conversion of astrocytes into neurons. (A) Schematic diagram showing virus injection sites and experimental design. (B) Representative images showing inversely correlated expression of GFP (green) and NeuroD1 (ND1, red). dpi, days post-virus injection. Scale bars, 50 μm. (C) Representative confocal images showing the relationship between GFP (green), ND1 (red), GFAP (violet), NeuN (cyan) staining. White arrowheads point to GFP+GFAP+ NeuN-astrocytes and red arrowheads point to GFP+NeuN+GFAP-neurons. Scale bars, 10 μm. (D,E) Quantification of GFP fluorescence and NeuroD1 intensity in mouse cortices injected with different AAV doses of NeuroD1. Values are shown as mean ± SD. N = 4 per group. (F) Quantification of the percentage of GFP+NeuN+ cells among total GFP+ cells in mouse cortices expressing different levels of NeuroD1. Values are shown as mean ± SD. N = 4 per group.

### Enhancing NeuroD1 expression with CMV enhancer increases conversion efficiency

Since increasing AAV dose might result in higher leakage, we designed a different approach to increase NeuroD1 expression by inserting a CMV enhancer in front of the GFAP promoter (GFAPCE::NeuroD1) and evaluated its effect on astrocyte conversion. We performed side-by-side comparisons between GFAPCE::NeuroD1 and GFAP104::NeuroD1, with the addition of GFAP104::GFP to label astrocytes (Fig. 3A). At 4 dpi, GFAP104::NeuroD1 showed very weak NeuroD1 expression and no significant effect on the GFP-labeled astrocytes (Fig. 3B); in contrast, GFAPCE::NeuroD1 showed much stronger NeuroD1 expression and a few GFP-labeled cells had lost astrocytic processes (Fig. 3C). By 30 dpi, the majority of GFP-labeled cells in the GFAP104::NeuroD1 group remained typical astrocytic morphology (Fig. 3D), but in the GFAPCE::NeuroD1 group, the majority of GFP-labeled cells changed into neuronal morphology with clear apical dendrites (Fig. 3E, white arrow). A series of immunostaining revealed that at 30 dpi, GFP-labeled cells in the GFAP104::NeuroD1 group were mostly colocalized with astrocytic marker GFAP, whereas the majority of GFP-labeled cells in the GFAPCE::NeuroD1 group were colocalized with neuronal marker NeuN (Fig. 3D-F). Quantitative analyses revealed that at 30 dpi, the conversion rate in the GFAPCE::NeuroD1 group (~70%) was much higher than that in the GFAP104::NeuroD1 group (~20%) (Fig. 3G), which correlated with higher NeuroD1 expression level in the GFAPCE::NeuroD1 group (Fig. 3H). Together, these results suggest that enhancing NeuroD1 expression level is critical to increase the efficiency of astrocyte-to-neuron conversion.

**Figure 3.**
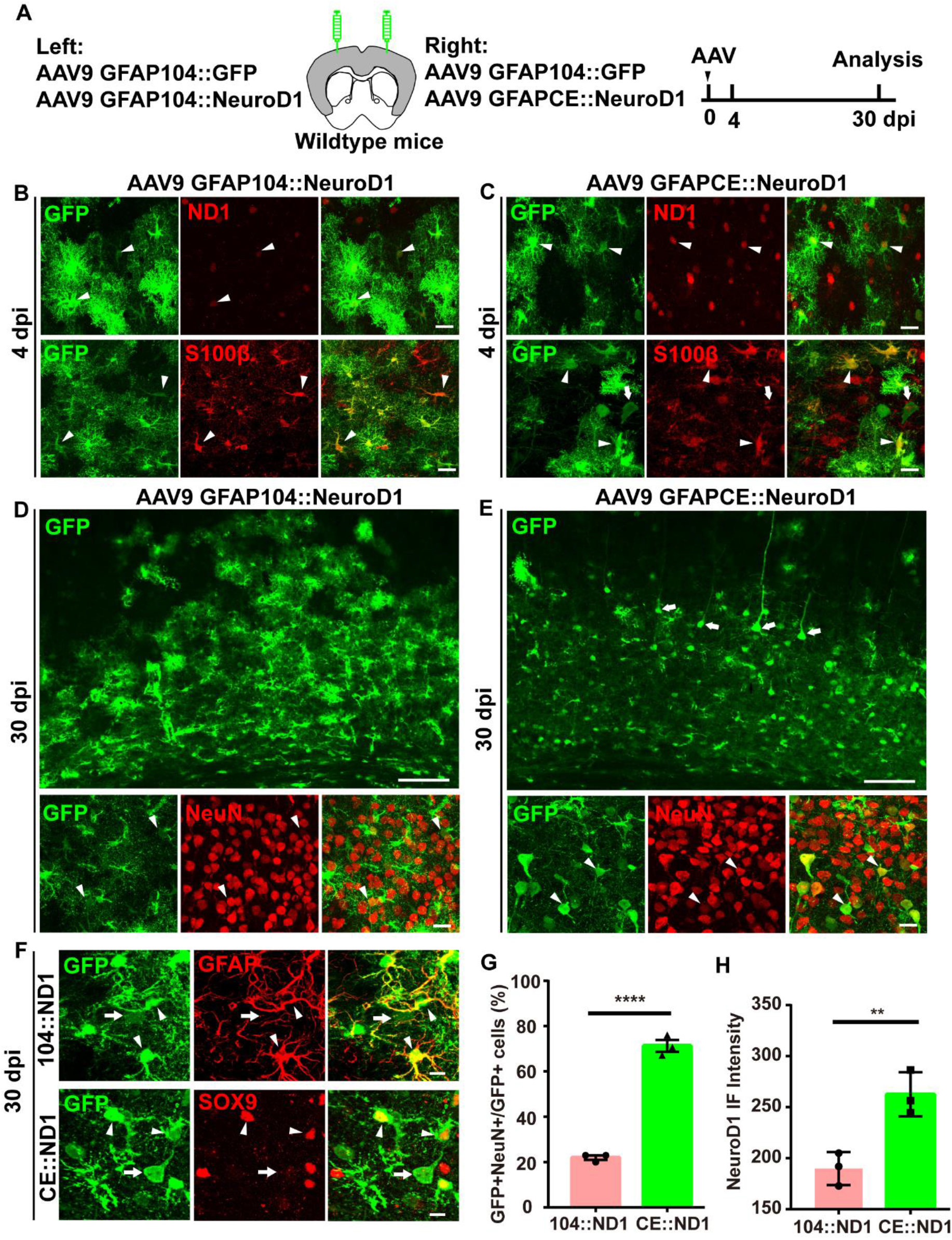
Enhancing NeuroD1 expression with CMV enhancer increases astrocyte-to-neuron conversion efficiency. (A) Schematic diagram illustrating experimental design. (B-C) Representative images showing co-immunostaining of GFP, NeuroD1 (ND1, red) and astrocytic marker S100β (red) at 4 days after AAV injection (dpi). Most of the GFP+ cells expressed both NeuroD1 and the astrocytic marker S100β at 4 dpi. Arrowheads point to GFP+NeuroD1+ or GFP+S100β+ co-labeled cells. Scale bars, 10 μm. (D-F) Representative images showing side-by-side comparisons between GFAP104::NeuroD1 and GFAPCE::NeuroD1 in inducing AtN conversion at 30 dpi. Most of the GFP+ cells in the GFAP104::NeuroD1 group showed astrocytic morphology and co-localized with astrocytic marker GFAP (D, F), whereas Most of the GFP+ cells in the GFAPCE::NeuroD1 group showed neuronal morphology with long apical dendrites and co-localized with neuronal marker NeuN (E-F). Scale bars, 100 μm for D-E top panels; 10 μm for D-E bottom panels and F. (G-H) Quantification of the conversion efficiency (G; GFP+NeuN+ cells among total GFP+ cells) and NeuroD1 expression level (H) in mouse cortices infected by GFAP104::NeuroD1 or GFAPCE::NeuroD1. Values are shown as mean ± SD. N = 3 per group. Student’s *t*-test, **p<0.01, ****p<0.0001.

### Astrocyte conversion in lineage-tracing mice

After understanding the effect of NeuroD1 expression on astrocyte conversion, we started to solve the puzzle why some labs including ours can convert lineage-traced astrocytes into neurons (Leib et al., 2022; Xiang et al., 2021), whereas some others cannot (Wang et al., 2021). We injected control AAV Flex-GFP or testing AAV Flex-NeuroD1-GFP into the astrocyte-specific transgenic mice (Aldh1l1-CreERT2) to investigate AtN conversion in lineage-traced cortical astrocytes (Fig. 4A). We found that injecting control AAV Flex-GFP labeled >90% of the cortical astrocytes (S100b+) in the lineage-traced mice, with <10% neuronal leakage by 60 dpi (Fig. 4B). Similarly, injecting AAV Flex-NeuroD1-GFP into the Aldh1l1-CreERT2 mouse cortex initially also labeled most astrocytes in green (Fig. 4C, 6 dpi); however, by 60 dpi, many GFP-traced astrocytes changed into neuronal morphology with clear apical dendrites (Fig. 4D, red arrow). Immunostaining results revealed that at 6 dpi after injecting Flex-NeuroD1-GFP, most of the GFAP+ astrocytes were labeled by GFP, and NeuroD1 expression was also clearly detected inside GFAP+ astrocytes (Fig. 4E). At 60 dpi after injecting Flex-NeuroD1-GFP, while many GFP-labeled cells remained astrocytic morphology and colocalized with astrocytic marker S100b, a significant portion of the GFP-traced astrocytes had turned into neurons and colocalized with neuronal marker NeuN (Fig. 4E-G, red arrows). Quantitative analyses revealed that after injecting Flex-NeuroD1-GFP into astrocytic lineage-tracing Aldh1l1-CreERT2 mice, there were >15% astrocyte-converted NeuN+ neurons at 30 dpi, and >25% astrocyte-converted NeuN+ neurons at 60 dpi (Fig. 4H). In addition, there were ~10% of NeuroD1-GFP-infected astrocytes changed into an intermediate stage that had lost astrocytic marker S100b but did not acquire neuronal marker NeuN at 60 dpi (Fig. 4H, NeuN-S100b-), suggesting that many astrocytes were still in the process of converting. Together, these results demonstrate that lineage-traced astrocytes in Aldh1l1-CreERT2 mice can be converted into neurons, but their conversion efficiency appears to be much lower than that in WT mice, suggesting higher conversion barrier in these lineage-traced astrocytes.

**Figure 4.**
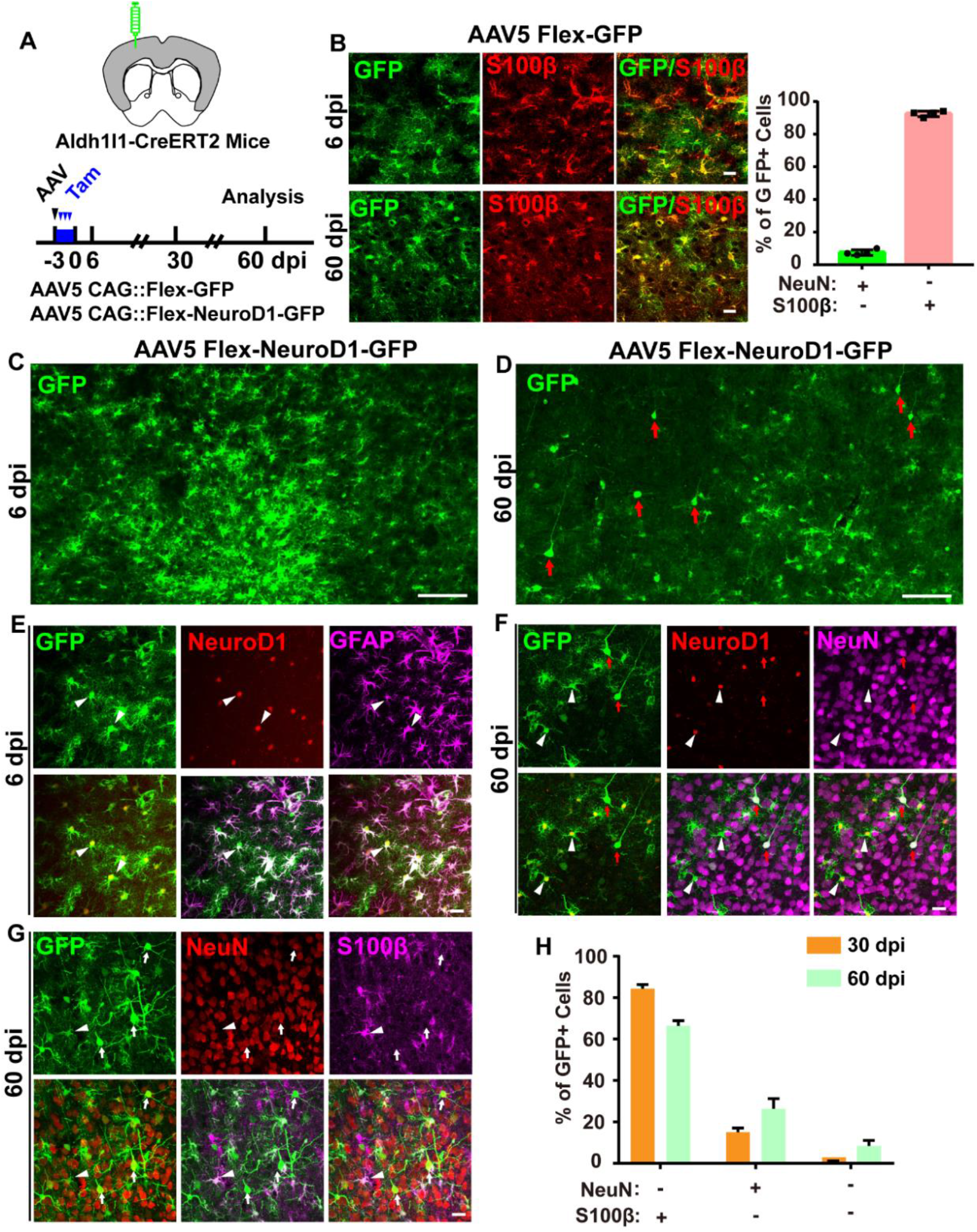
Astrocyte-to-neuron conversion in astrocytic lineage-tracing mice. (A) Schematic diagram showing experimental design. (B) Specific labeling of astrocytes in Aldh1l1-CreERT2 lineage-tracing mice. Representative images (left panels) showing co-localization of GFP and astrocytic marker S100β (red) at 6 and 60 dpi in the control group (AAV5 Flex-GFP). Scale bars, 10 μm. Quantification (right bar graph) of the percentage of NeuN+ or S100β+ cells within GFP+ cells at 60 dpi. >90% of GFP-expressing cells were S100β+ astrocytes. Values are shown as mean ± SD. N = 4 per group. (C-D) Time-dependent conversion of lineage-traced astrocytes into neurons. Representative images showing the comparison between the GFP+ cells at 6 dpi (C) versus 60 dpi (D). Most of the GFP+ cells showing astrocytic morphology at 6 dpi; but some GFP+ cells showing neuronal morphology with long apical dendrites at 60 dpi. Red arrows point to GFP+ cells with typical neuronal morphology. Scale bars, 100 μm. (E-F) Representative images showing immunostaining of GFP (green), NeuroD1 (red), astrocytic marker GFAP (violet, E) or neuronal marker NeuN (violet, F) at 6 (E) and 60 (F) dpi. Most of the Flex-ND1-GFP-infected cells expressed NeuroD1 and were colocalized with astrocytic marker GFAP at 6 dpi (E). By 60 dpi, some Flex-ND1-GFP-infected cells obtained neuronal morphology and were colocalized with neuronal marker NeuN (F). White arrowheads point to GFP+ cells expressing NeuroD1, while red arrows point to GFP+ cells expressing neuronal marker NeuN. Scale bars, 10 μm. (G-H) Assessing conversion efficiency in astrocytic lineage-tracing mice. Representative images showing GFP, S100β (violet) and NeuN (red) at 60 dpi (G). Arrows points to GFP+NeuN+ neurons, while arrowheads point to GFP+S100β+ astrocytes. Scale bars, 10 μm. Quantification of the percentage of NeuN+ or S100+ or NeuN-S100-cells among GFP+ cells at 60 dpi. Note that even at 60 dpi, the conversion efficiency in lineage-traced astrocytes only reached over 20%, suggesting high conversion barrier in these lineage-traced astrocytes. Values are shown as mean ± SD. N = 3 per group.

### Converting lineage-traced astrocytes using CMV enhancer to increase NeuroD1 expression

We were alerted by the apparent lower conversion efficiency in the lineage-tracing Aldh1l1-CreERT2 mice compared to our previous results obtained in WT mice (Chen et al., 2020; Ge et al., 2020; Guo et al., 2014; Liu et al., 2020; Puls et al., 2020; Tang et al., 2021; Wu et al., 2020; Zhang et al., 2020; Zheng et al., 2022). To increase the conversion rate in lineage-traced astrocytes without increasing AAV titre, we employed AAV vector with CMV enhancer (GFAPCE::NeuroD1) engineered to increase NeuroD1 expression in astrocytes as shown in Figure 3. In this set of experiments, we crossed Aldh1l1-CreERT2 mice with Ai14 mice so that most of the astrocytes were labeled by red fluorescent protein tdTomato following the administration of tamoxifen (Fig. 5A-B). After injecting AAV5 GFAPCE::NeuroD1 into one side of the mouse cortex, we found that while some tdTomato-labeled cells were still GFAP+ astrocytes, some other tdTomato-labeled cells showed reduced tdTomato signal and colocalized with NeuN signal at 30 dpi (Fig. 5C-D). By 60 dpi following infection of GFAPCE::NeuroD1, we detected many tdTomato-labeled cells in typical pyramidal neuron shape and colocalized with NeuN (Fig. 5E-H), indicating that the lineage-traced tdTomato-labeled astrocytes had been successfully converted into neurons. Quantitative analyses revealed that at 60 dpi after injecting AAV GFAPCE::NeuroD1 into one side of the cortex, there were ~40% of tdTomato-traced astrocytes converted into NeuN+ neurons (NeuN+tdTomato+/tdTomato+), and ~40% of tdTomato-traced astrocytes remained GFAP+ astrocytes (GFAP+tdTomato+/tdTomato+) (Fig. 5I, gray bar). The other 20% tdTomato+ cells did not show GFAP signal nor NeuN signal (GFAP-NeuN-/tdTomato+), likely in a transitional stage during AtN conversion process. In contrast, in the control side where no NeuroD1 was injected, >90% tdTomato+ cells were GFAP+ astrocytes (Fig. 5I, white bar). Similarly, when counting the total number of NeuN+tdTomato+ neurons, there were 13.3 ±1.1 tdTomato+ neurons/mm^2^ in the control side, suggesting that even such Aldh1l1-CreERT2 mice still has neuronal leakage. Nevertheless, in the NeuroD1-injected side, there were 235.1 ± 22.4 tdTomato+ neurons/mm^2^. Clearly, GFAPCE::NeuroD1 has converted tdTomato-traced astrocytes into tdTomato+ neurons.

**Figure 5.**
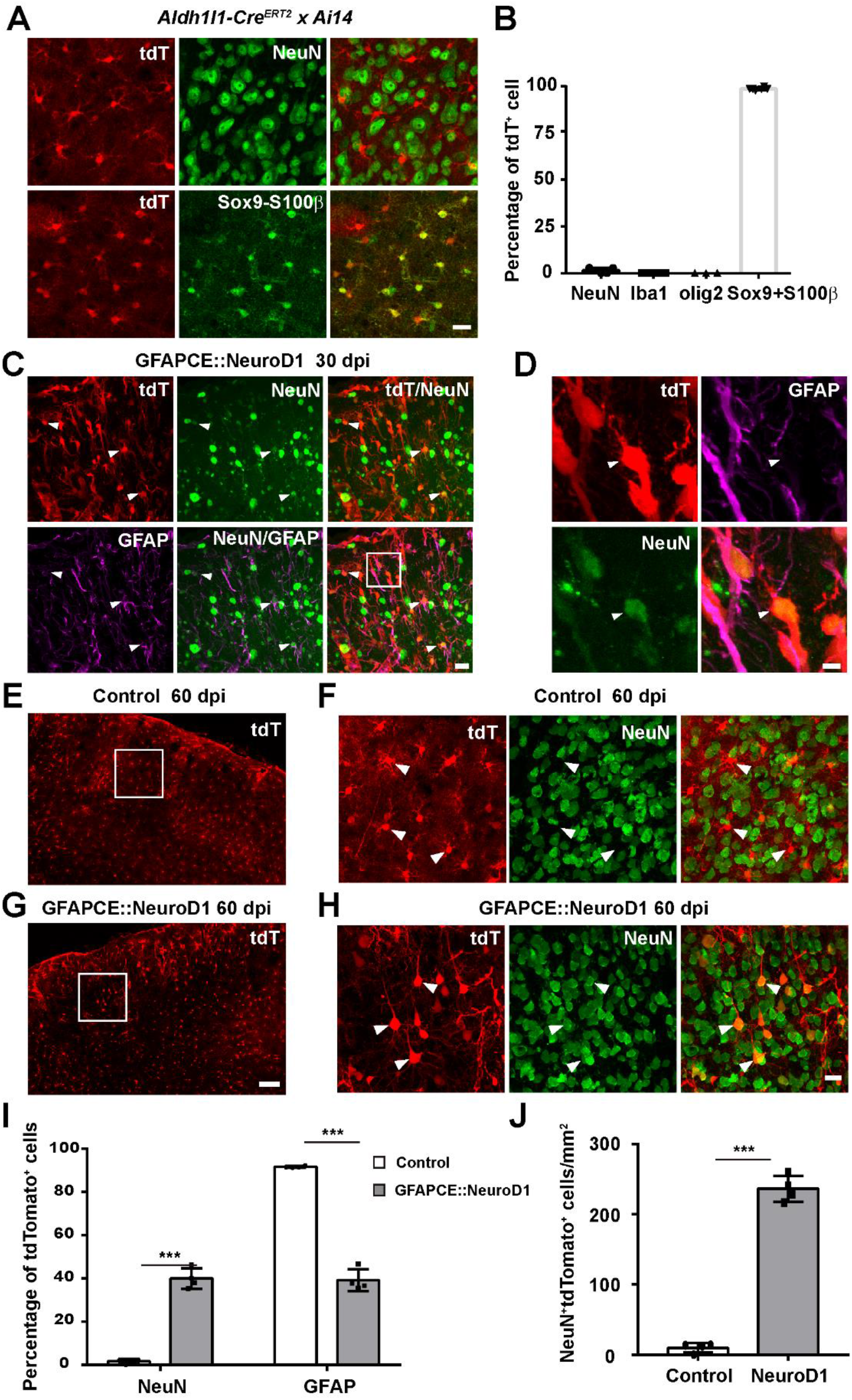
Enhancing NeuroD1 expression with CMV enhancer converts lineage-traced tdTomato^+^ astrocytes into tdTomato^+^ mature neurons. (A) Representative images showing immunostaining of tdTomato (red), neuronal marker NeuN (green) and astrocytic markers Sox9+S100β (green) in the Aldh1l1-Cre^ERT2^ X Ai14 mouse cortex. Note that the vast majority of tdTomato^+^ cells were Sox+S100β+ astrocytes in Aldh1l1-Cre^ERT2^ X Ai14 mouse brains. Scale bar, 20 μm. (B) Quantification of the percentage of NeuN+ or Iba1+ or oligo2+ or Sox9+S100β+ cells among tdTomato+ cells in Aldh1l1-Cre^ERT2^ X Ai14 mouse cortex. Values are shown as mean ± SD. N = 3 per group. (C-D) Representative images showing immunostaining of tdTomato (red), NeuN (green) and GFAP (violet) in Aldh1l1-Cre^ERT2^ X Ai14 mouse cortex at 30 days after injecting GFAPCE::NeuroD1 into one side cortex. Note that in NeuroD1-injected areas, some tdTomato+ cells had already changed their morphology and lost GFAP signal but acquired NeuN signal at 30 dpi (D, arrowhead). Scale bar, 20 μm for C, 5 μm for D. (E-H) Comparison between tdTomato^+^ cells without (E-F) or with GFAPCE::NeuroD1 infection (G-H). Representative images showing tdTomato (red) and NeuN (green) staining in Aldh1l1-Cre^ERT2^ X Ai14 mouse cortex at 60 dpi. Note that the tdTomato^+^ cells in control side (E-F, no-NeuroD1 infection) were rarely NeuN+, but many tdTomato^+^ cells in NeuroD1-infected areas were NeuN+ neurons with clear apical dendrites (G-H). (F) and (H) are high-magnification images enlarged from the regions in the white boxes in (E) and (G), correspondingly. Scale bar, 100 μm (E, G), 10 μm (F, H). (I-J) Quantification of the percentage of NeuN+ neurons or GFAP+ astrocytes among total tdTomato+ cells (I) or the actual number of tdTomato+NeuN+ neurons (J) at 60 dpi. Note that compared to the control side, GFAP::NeuroD1-infected side showed a significant decrease of tdTomato+GFAP+ astrocytes, which was accompanied with a significant increase of tdTomato+NeuN+ neurons. Values are shown as mean ± SD. N = 4 per group. Student’s *t*-test, ***p<0.001.

### Two-photon live imaging of astrocyte-to-neuron conversion

To unambiguously demonstrate astrocyte-to-neuron conversion, we further employed two-photon microscopy to lively visualize the conversion process in the mouse cortex. When we injected control AAV9 GFAP::GFP into the mouse cortex, the GFP-labeled cells displayed typical astrocytic morphology (Fig. 6A), which were also immunostained positive for astrocytic marker GFAP after fixation (Fig. 6D). In contrast, after injecting AAV9 GFAP::NeuroD1-GFP into the mouse cortex, the GFP-labeled cells initially also displayed astrocytic morphology but gradually changed into neuronal morphology within 3 weeks (Fig. 6C, n = 10 cells from 4 mice). Note that between the two GFP-labeled cells, there was a blood vessel (Fig. 6C, 14 dpi, white arrow) that could be used as a landmark to unambiguously identify the two astrocytes undergoing conversion process. After two-photon live imaging, we performed post-fixation immunostaining and found that the majority of GFAP::NeuroD1-GFP-infected cells were NeuN+ neurons (Fig. 6E). Besides AAV, we also employed retrovirus that only infects dividing glial cells to monitor conversion process under two-photon microscope. Indeed, we observed glial cell division infected by control retrovirus CAG::GFP in the mouse cortex (Fig. 6B). After injecting retrovirus CAG::NeuroD1-GFP into the mouse cortex, the NeuroD1-infected cells showed rapid morphological change within 6 days—changing from glial cells into neuron-like cells with long axon-like processes (Fig. 6F, n = 11 cells from 3 mice). After live imaging, post-fixation immunostaining also confirmed that the retrovirus NeuroD1-GFP-infected cells were NeuN+ neurons (Fig. 6G). Moreover, we performed two-photon live imaging in astrocytic lineage-tracing mice (Aldh1l1-CreERT2 x Ai14) and injected AAV5 GFAPCE::NeuroD1 into the mouse cortex following the administration of tamoxifen (Fig. 6H). Clearly, under two-photon microscope, the tdTomato-labeled astrocytes gradually changed into tdTomato-labeled neurons with long apical dendrites (Fig. 6H, see also Suppl. Movie 1 and 2). Together, these two-photon microscopic experiments unambiguously captured the astrocyte-to-neuron conversion process lively in the mouse brain.

**Figure 6.**
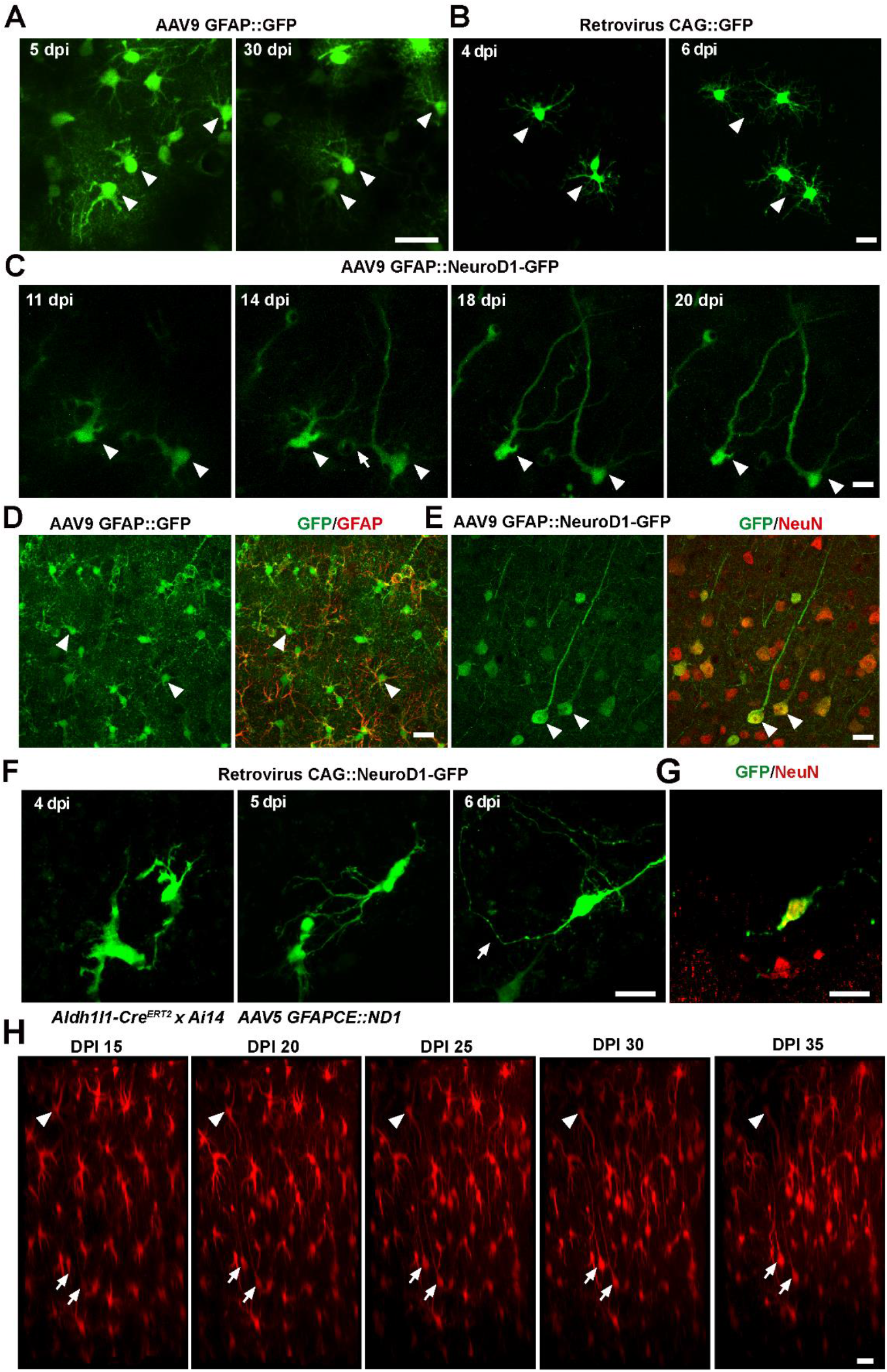
*In vivo* two-photon live imaging of NeuroD1-induced astrocyte-to-neuron conversion. (A,B) Representative two-photon images of GFP-labeled astrocytes infected by control AAV9 GFAP::GFP (A) or control retrovirus CAG::GFP (B). Note that the control AAV GFAP::GFP virus-infected astrocytes did not change their astrocytic morphology and stayed in the same position over 30 days (A). Control retrovirus CAG::GFP-infected cells showing cell division over 6 days (B). Scale bar, 20 μm (A), 10 μm (B). (C) Two-photon time-lapse images illustrating morphological changes of GFP-labeled astrocytes within 3 weeks after infected by AAV9 GFAP::NeuroD1-P2A-GFP. The NeuroD1-GFP-infected cells (arrowhead) initially showed astrocytic morphology but gradually changed into neuronal morphology with long apical dendrites. Scale bar, 20 μm. (D) Representative images showing control GFAP::GFP-infected cells (green) were immunopositive for astrocytic marker GFAP. Scale bar, 20 μm. (E) Representative images showing GFAP::NeuroD1-GFP-infected cells (green) were NeuN-positive neurons at 30 dpi. Scale bar, 20 μm. (F) Representative two-photon images showing rapid astrocyte-to-neuron conversion after infected by retrovirus CAG::NeuroD1-GFP. Note that by 6 DPI, a converted neuron already exhibited a long and curved axon (white arrow). Scale bar, 20 μm. (G) Representative image showing neuronal marker NeuN staining of the newly converted cell in F (white arrow), confirming its neuronal identity. Scale bar, 20 μm. (H) Representative two-photon live imaging of tdTomato+ lineage-traced astrocytes after AAV5 GFAPCE::NeuroD1 infection. Many tdTomato+ cells gradually changed from astrocytic morphology at 15 dpi to typical neuronal morphology with clear apical dendrites by 35 dpi after GFAPCE::NeuroD1-infection. Scale bar, 20 μm.

### NeuroD1-induced astrocyte conversion in the striatum

So far we have provided ample evidence that different NeuroD1 vectors can convert astrocytes into neurons in the mouse cortex with different genetic background. We have previously demonstrated that NeuroD1 mainly converts cortical astrocytes into glutamatergic pyramidal neurons (Chen et al., 2020; Ge et al., 2020; Tang et al., 2021; Zhang et al., 2020). We then asked a further question: if injecting NeuroD1 into the striatum, where most neurons are DARPP32+ GABAergic neurons, will NeuroD1 generate striatal neurons or cortical neurons? Of course, if NeuroD1 cannot convert astrocytes at all but only leaks into native neurons as claimed by Wang et al (2021), then we would expect that all neurons infected by NeuroD1 would be DARPP32+ GABAergic neurons. To test this possibility, we injected GFAP104::NeuroD1-GFP into the mouse striatum and performed a series of immunostaining at 4 dpi, 30 dpi and 60 dpi (Fig. 7A). At 4 dpi, essentially all cells infected by AAV9 GFAP104::NeuroD1-GFP were S100β+ astrocytes and rarely any NeuN+ neurons (Fig. 7B-C). More importantly, NeuroD1 immunostaining revealed that they were all colocalized with GFAP+ astrocytes (Fig. 7D). At 30 dpi, we found that GFAP104::NeuroD1-GFP-infected cells were a mixture of Sox9+ astrocytes and NeuN+ neurons (Fig. 7E-F). Quantitative analysis revealed that ~40% of NeuroD1-GFP-infected cells showed NeuN signal, but only <10% of the NeuroD1-GFP-infected cells showed striatal medium spiny neuron (MSN) marker DARPP32 (Fig. 7F-G). By 60 dpi, the majority of NeuroD1-GFP-infected cells had gained NeuN signal (Fig. 7H-I). NeuroD1 immunostaining also revealed their presence in neuron-like cells (Fig. 7I). Quantified results showed that >80% of NeuroD1-GFP-infected cells were NeuN+ neurons, but still only <30% showed DARPP32 signal (Fig. 7I-J). Thus, the fact that the majority of NeuroD1-converted neurons in the striatum are not native striatal neurons strongly argues against NeuroD1 leakage into preexisting neurons. Instead, these non-native neurons generated by NeuroD1 at 60 dpi can only be converted from those NeuroD1-expressing astrocytes at 4 dpi.

**Fig. 7.**
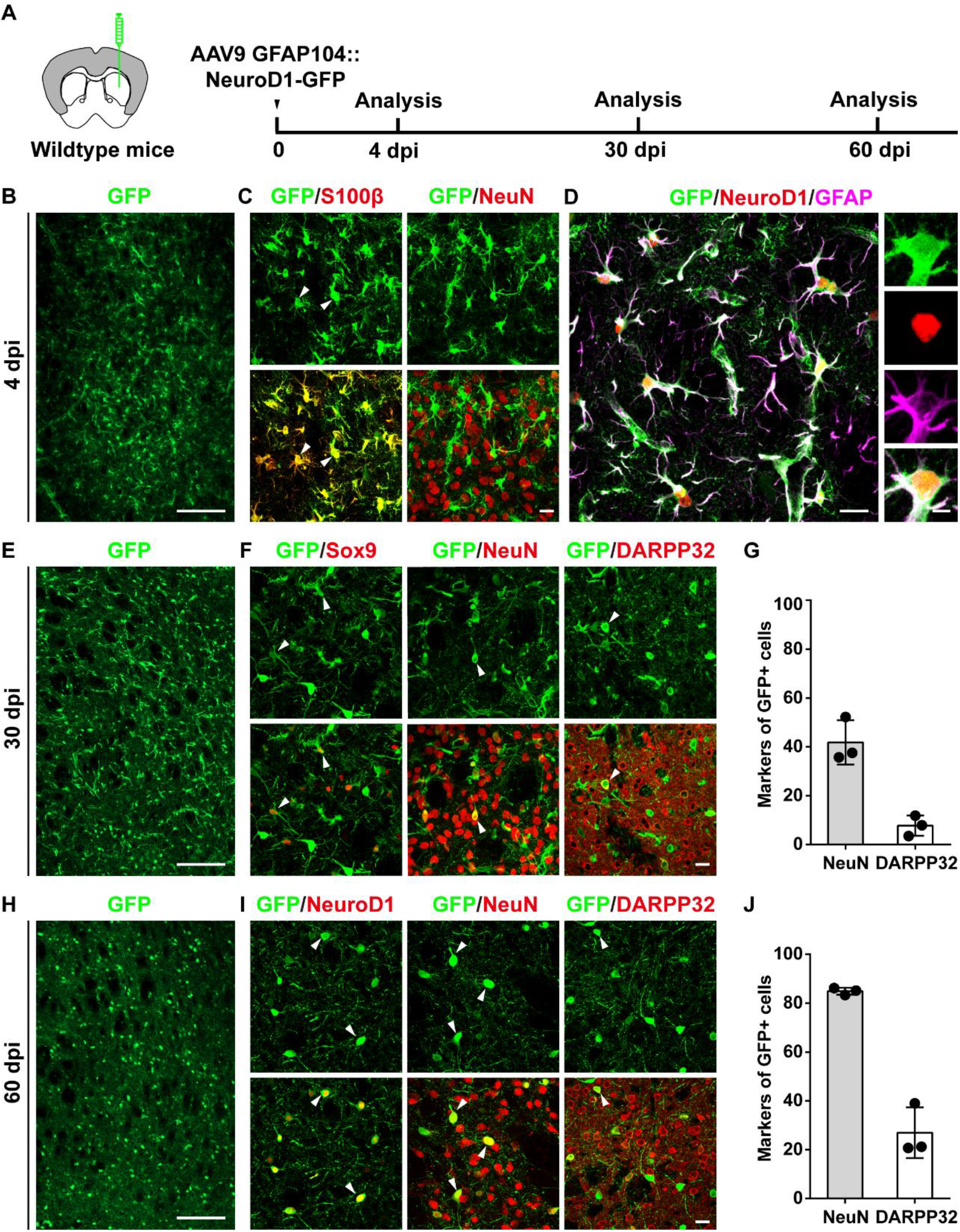
*In vivo* NeuroD1-indued striatal astrocyte-to-neuron conversion. A. Experimental design. B. Representative image showing wide infected area by AAV9 GFAP104::NeuroD1-GFP (green) in the striatum at 4 dpi. Scale bar: 200 μm. C. Confocal images showing that at 4 dpi most NeuroD1-GFP-infected cells (green) were co-stained with astrocytic marker S100β (red, left column) but rarely with neuronal marker NeuN (red, right column). Arrowheads indicate GFP and S100β colocalized cells. Scale bar, 20 μm. D. Confocal images showing clear NeuroD1 expression (red) in GFAP+ astrocytes (magenta). Scale bar, 20 μm and 5 μm. E. Low magnification image showing AAV9 GFAP104::NeuroD1-GFP (green) infected striatal area at 30 dpi. Scale bar, 200 μm. F. Confocal images showing that NeuroD1-GFP-infected cells (green) were partially co-stained with astrocytic marker Sox9 (red, left column) or neuronal marker NeuN (red, middle column), or with medium spiny neuron marker DARPP32 (red, right column). Arrowheads indicate some colocalized cells. Scale bar, 20 μm. G. Quantification of conversion rate at 30 days after NeuroD1 infection in the striatum. Note that although about 41% (41.8 ± 9.1%) of NeuroD1-GFP-infected astrocytes were converted into NeuN+ neurons, the proportion of converted MSNs was limited (7.7 ± 4.2%). N = 3 animals. H. Representative image showing NeuroD1-GFP (green) infected wide striatal area at 60 dpi. Scale bar, 200 μm. I. Confocal images showing that by 60 dpi, NeuroD1-GFP-infected cells (green) still expressed NeuroD1 (red, left column) but this time they were converted into NeuN+ neurons (red, middle column), with some also immunopositive for DARPP32 (red, right column). Arrowheads indicate some colocalized cells. Scale bar, 20 μm. J. Quantified date showing the conversion rate at 60 dpi reaching over 80% (84.9 ± 1.4%), but only a small portion being DARPP32+ MSNs (27.0 ± 10.4%). N = 3 animals.

## DISCUSSION

In this study, we demonstrate with multiple lines of evidence that astrocytes from both WT or lineage-tracing mice can be converted into neurons. However, astrocytes in WT mice are much easier to convert than those in lineage-tracing mice, and the conversion efficiency is much higher in WT mice than that in lineage-tracing mice. Therefore, lineage-tracing mice pose higher conversion barrier that must be overcome when conducting AtN conversion studies. We further demonstrate here that one potential solution to overcome the higher conversion barrier is to use enhancer to increase NeuroD1 expression in the lineage-traced astrocytes. Under two-photon microscope, we can also lively monitor the AtN conversion process in the mouse cortex, including the lineage-traced astrocytes. Thus, these results demonstrate from different angles that NeuroD1 can convert astrocytes into neurons *in situ*.

### High AAV titre causing high neuronal leakage

Injecting viruses into a healthy brain has been widely used as an important research tool to understand brain circuit formation and functions. However, how much virus can be administered into a healthy brain is often a neglected topic. If injecting a large amount of viruses into a healthy brain, viral infection and their transgene expression will certainly trigger anti-viral responses in brain cells that may result in altered brain functions. The more virus injected, the more distorted functions one may get from the viral infected brain cells. Then, how do we know the right amount of viruses to be injected into a healthy brain or an injured brain in order to avoid serious side effects that may compromise our data interpretation? The right answer is to do a series of dose-finding studies in order to identify a right dose that will minimize side effects. Our dose-finding studies demonstrate that AAV vectors injected into healthy mouse brains shall be kept in a titre range of 10^11^ – 10^12^ GC/ml, assuming the volume being kept at 1-2 μl as commonly used for most mouse brain studies. When AAV titre reaches 10^13^ GC/ml, AAV vector GFAP::GFP starts to express GFP significantly in neuronal cells, breaking down the restriction of GFAP promoter in astrocytes and leading to so-called “neuronal leakage”. Such neuronal leakage is likely due to high dosage of AAV causing damages to both neurons and astrocytes as reported before (Ortinski et al., 2010). It has also been reported that damaged neurons may have low level of GFAP expression, indicating that GFAP promoter can be partially active in damaged neurons (Zwirner et al., 2021). Neuronal leakage may also be a by-product of cellular immune reaction. When massive AAV viruses attack brain cells including neurons and astrocytes, they may trigger strong anti-viral responses in brain cells (Hudry and Vandenberghe, 2019). However, if one injects too much AAV such as high titre of 10^13^ GC/ml at 1 μl, it can be overwhelming for brain cells to deal with and some brain cells may get injured or even die in the battle against the viruses. It is likely those injured neurons that will show leakage after being attacked by a large amount of viruses.

Here, we want to emphasize the side effects of high titre instead of total dose because high titre viruses in low volume may cause even more damage than low titre viruses in high volume. After all, when injected at high titre, each brain cell at the injecting site will be surrounded by much more viral particles than in low titre case. Given our dose-finding studies that high titre AAV of 10^13^ GC/ml at 1 μl causing significant neuronal leakage in the mouse brain, we suggest that previous studies using high titre AAV of 10^13^ GC/ml or above such as those by Wang et. (Wang et al., 2021), Mattugini et al. (Mattugini et al., 2019), and Zhou et al. (Zhou et al., 2020) be reevaluated with lower titre in the range of 10^11^ – 10^12^ GC/ml (1 μl) in order to validate their conclusions.

#### Conversion versus leakage

One common misunderstanding in the AtN conversion field is that if neuronal leakage occurs, then AtN conversion is in doubt. Such one-way thinking is a misunderstanding of the relationship between AtN conversion and neuronal leakage. In fact, AtN conversion and neuronal leakage are two relatively independent events that often occur simultaneously in one conversion study. As discussed above, neuronal leakage is due to some AAV vectors (such as GFAP::GFP) directly express their transgene (such as GFP) in neurons besides in astrocytes where they should express. However, neuronal leakage itself does not mean the astrocytes expressing neural transcription factors cannot be converted into neurons. Neuronal leakage and astrocyte conversion are not mutually exclusive. Even at very high AAV titre of 10^13^ GC/ml, we show that the leakage rate is 50-60% but not 100%. In Wang et al. (Wang et al., 2021), they reported a neuronal leakage rate of 37.5% in NeuroD1 group by using retrograde tracing technique, they did not explain where the other 62.5% neurons came from. Clearly, one possibility is that the other 62.5% neurons were converted from NeuroD1-expressing astrocytes in WT mice. We have repeatedly observed that in NeuroD1-mediated astrocyte conversion, NeuroD1 expression was initially detected in GFAP+ astrocytes within the first week of viral infection, but later detected in NeuN+ neurons after several weeks. If it is not because those NeuroD1-expressing astrocytes being converted into NeuroD1-expressing neurons, how can one explain the fact that NeuroD1 jumps from astrocytic nucleus to neuronal nucleus? In fact, when NeuroD1 is expressed in astrocytes, nothing can stop its function in converting astrocytes into neurons, so long as the expression level of NeuroD1 is high enough to overcome the conversion barrier. This has been shown clearly in our recent RNA-seq analyses demonstrating that NeuroD1 expression in astrocytes downregulates glial genes and upregulates neuronal genes to turn the astrocytic transcriptome into neuronal transcriptome within 2 weeks (Ma et al., 2022).

Neuronal leakage is a default phenomenon when one injects billions of AAV particles into a normal healthy brain. Why convert astrocytes into neurons in a healthy brain? AtN conversion is meant to generate new neurons in conditions with significant neuronal loss. If someone insists to inject a huge number of viruses into a healthy brain, then be prepared to see leaked neurons due to viral injury. The real question to ask for any AtN conversion study is “how many neurons are converted and how many neurons are leaked?” Obviously, the goal is to increase the proportion of converted neurons (such as using an enhancer) and decrease the proportion of leaked neurons (such as lower the titre). Since both AAV and lentivirus can infect neurons and astrocytes, it is almost impossible to keep neuronal leakage down to 0%. Typically, we recommend to keep the leaked neurons below 10% among the mixture of leaked and converted neurons altogether.

### Wildtype mice versus lineage-tracing mice

Lineage tracing mice such as Aldh1l1-CreERT2 mice were created to pre-label astrocytes in order to trace where they are and where they come from. They are not created for reprogramming purpose. If we want to convert the lineage-traced astrocytes into neurons, we must ask whether they are the same as WT astrocytes when infected by the same virus expressing the same transcription factor(s). Without such strict side-by-side comparison studies, it is pre-mature to claim any golden standard for reprogramming purpose if we don’t even understand these lineage-traced astrocytes in terms of conversion capability. Here, we demonstrate that NeuroD1 converts WT astrocytes into neurons with high efficiency, but in Aldh1l1-CreERT2 mice the conversion rate is much lower in lineage-traced astrocytes. These results suggest that the lineage-traced astrocytes may have higher conversion barrier than the WT astrocytes. While precise mechanisms behind such high barrier in lineage-traced astrocytes warrant further study, one possibility is that lineage-traced astrocytes require Cre-LoxP-mediated DNA recombination, which involves DNA cutting and annealing. For any healthy cell, DNA cutting and annealing is a kind of DNA damage which may trigger massive cellular responses including DNA methylation and other epigenetic modifications (Karakaidos et al., 2020; Livingston et al., 2020; Loonstra et al., 2001). Given the fact that the lineage-traced astrocytes behave very differently from WT astrocytes in terms of conversion, we strongly advise the entire AtN conversion field to be extra cautious when using lineage-traced astrocytes for reprogramming purpose. While lineage-traced astrocytes are a valid tool to study where newly generated neurons come from, we shall be fully aware of their high conversion barrier and try every effort possible to overcome such barrier in order to draw solid conclusions. If no effort made at all to reduce the high conversion barrier or to enhance the expression level of converting factors, it will not be a surprise that one may not be easy to find converted neurons. It is not because the astrocytes cannot be converted; it is rather because of a lack of understanding of these linage-traced astrocytes and insufficient driving power to convert them.

### Healthy brain versus injured brain

Before conducting any AtN conversion study, one shall ask “what is the purpose to do AtN conversion study?” Under what circumstance shall we convert an astrocyte into a neuron? A healthy brain is composed of neurons and glial cells in a delicate balance. If we inject viruses into a healthy brain to convert astrocytes into neurons, we are surely breaking down the delicate balance between neurons and astrocytes and causing injuries to the healthy brain. The degree of injury depends on the amount of viruses injected into the healthy brain. Therefore, it is never a wise idea to inject viruses into a healthy brain for cell conversion purpose, particularly when one injects a reckless amount of viruses into a healthy brain.

The purpose of AtN conversion should be to regenerate functional new neurons in injured or diseased brains, not healthy brains. Healthy brain does not need new neurons at all. Human brain injury often involves millions or even billions of neuronal loss. Over the past decades, there has been little progress in regenerating millions let alone billions of functional new neurons in mammalian brains to treat severe brain disorders such as stroke and Alzheimer’s disease. It is under such premise that AtN conversion technology is invented to regenerate functional new neurons in large quantity to repair damaged brains. The definition of large quantity here should not be measured by thousands but by millions at least. Because human brain has 86 billions of neurons, generating thousands of new neurons is not meaningful in terms of brain repair. In the case of stroke or Alzheimer’s disease, it may require tens or hundreds of millions of new neurons to have real meaningful functional recovery. Fortunately, every neuron is surrounded by a number of glial cells. Even better, glial cells can divide to replenish themselves, making them a youth fountain to generate new neurons without the worry of glial depletion. Our previous studies have demonstrated that after converting NeuroD1-infected astrocytes into neurons, the remaining non-infected astrocytes can divide to repopulate themselves (Zhang et al., 2020), suggesting that AtN conversion has the potential to repeatedly convert astrocytes into neurons in injured or diseased brains.

### Mechanistic study versus therapeutic investigation

Last but not least, when we conduct AtN conversion studies, we must be clear whether we want to solve a mechanistic problem or we want to solve a therapeutic problem. The experimental strategy can be different. For example, if we want to conduct a mechanistic study, and particularly want to investigate conversion mechanisms in a healthy brain, we shall inject very low titre viruses into a healthy brain, such as in the order of magnitude of ~10^11^ GC/ml AAV, to have minimal neuronal leakage. On the other hand, if we want to investigate therapeutic effect of AtN conversion and the purpose is to generate as many new neurons as possible, then we should use higher AAV titre such as in the order of magnitude of ~10^12^ GC/ml AAV. Of course, one shall bear in mind that 10^12^ GC/ml AAV will certainly result in higher neuronal leakage than low titre AAV at 10^11^ GC/ml. The question then becomes whether such leakage rate is acceptable for therapeutic purpose? Or in other words, what happens to the leaked neurons? In the case of NeuroD1, we did not observe any adversary effect of NeuroD1 during our stroke study (Chen et al., 2020) or glial scar reversal study (Zhang et al., 2020). After all, NeuroD1 is expressed in adult mammalian brains (Gao et al., 2009; Kuwabara et al., 2009), and we have detected low level of NeuroD1 expression in both rodents and non-human primate brains. Therefore, when one designs conversion studies, the selection of proper viral dosage (including titre and volume) should be dependent on the purpose of mechanistic study or therapeutic investigation. Ultimately, we suggest that every investigator should perform dose-finding studies using their own viruses in order to find the proper viral dose for conversion studies.

### Conclusion and Perspective

If someone injects stem cells into a healthy human brain to generate new neurons, it will surely be thought unethical. Now, if someone injects viruses into a healthy mouse brain in order to generate new neurons, how shall we judge such action? We believe that converting resident glial cells into new neurons in a healthy brain will inevitably have negative impact on the preexisting neural circuits and cause unwanted side effects. We advocate that AtN conversion studies be conducted in neural injury or disease models. The advantage of using internal dividing glial cells, such as astrocytes, to directly regenerate functional new neurons is an economic way to make use of our internal cell bank for regenerative purpose, but only when neurons are already lost. Because astrocytes can divide and divide again *in situ*, it is possible to administer multiple rounds of conversion factors to regenerate multiple rounds of new neurons. Therefore, if one wants to regenerate millions or even billions of new neurons in patients’ brain, there is no better way to regenerate new neurons using the proliferative glial cells next to the dead neurons. Since AAV is current choice for most gene therapy vectors, the challenge is to find the right AAV serotype and right dose that will efficiently generate the right type of neurons in the right place.

To help the AtN conversion field moving in the right direction, here we try to set up a general standard that should be easy to follow for anyone who would like to conduct AtN conversion studies. To identify true neuronal conversion induced by transcription factors (TFs) and avoid high neuronal leakage, we propose the following standard tests as the measure of successful AtN conversion:

1. All conversion studies should be ideally performed in injury or disease models.
2. For any kind of viral injection into the brain, spinal cord or retina, a dose-finding study should be performed to identify the optimal dose for conversion. For AAV in particular, the titre should be kept in the range of 10^11^ – 10^12^ GC/ml with a small volume (1-2 μl in mouse brain).
3. TF expression in astrocytes must be detected in early conversion stage such as within the first week.
4. For lineage-traced astrocytes, strong promoters and enhancers should be used to enhance TF expression at a high level for a sufficient time in order to trigger successful conversion.
5. Perform time course study to closely monitor astrocytic morphological changes as well as gradual loss of astrocytic markers in the TF-expressing astrocytes.
6. Identify intermediate stage in-between astrocytes and neurons during AtN conversion process that may show markers of both astrocytes (GFAP/S100b/Sox9) and neurons (NeuN), or neither astrocytic nor neuronal markers.
7. The ultimate success of AtN conversion is the regeneration of functional new neurons in injured or diseased areas, where TFs have been initially detected in astrocytic nuclei but later detected in neuronal nuclei.
8. Success of AtN conversion also means that among TF-expressing cells, the proportion of astrocytes will gradually decrease, accompanied by simultaneous increase of neuronal proportion. Astrocytic proliferation during conversion process can be monitored by cell proliferation markers such as Ki67.

With the settlement of conversion versus leakage issue, we can now spend more time in conducting AtN conversion studies in injury and disease models to find real cure for millions of patients suffering from neurological disorders. After all, there are no Cre-LoXP-recombinated astrocytes in human patients.

## Materials and methods

### Animals

Experiments were conducted on wild-type C57BL/6J mice (Guangdong Yaokang Biotechnology Co., Ltd., China). Aldh1l1-CreER^T2^ transgenic mice (#031008) and Ai14 knock in mice (#007914) were purchased from Jackson Laboratory. The mice were housed in a standard 12 hours light and dark cycle with sufficient water and food. The experimental protocols were approved by the Laboratory Animal Ethics Committee of Jinan University, China (approval No. IACUC-20210406-03).

### Virus production

Retroviral vector CAG-GFP and CAG::NeuroD1-IRES-GFP were constructed and packaged as previously described (Guo et al., 2014) for the retrovirus experiment. The titre of retrovirus was about 1~5e7 TU/ml. Single-stranded adenovirus-associated viral vectors (GFAP::GFP, GFAP::ND1-P2A-GFP, Flex-GFP, Flex-ND1-P2A-GFP) were constructed as previously described (Chen et al., 2020). GFAP promoters used in the experiment included GFAP2.2 (length 2200 bp, gfa2 promoter), GFAP1.6 (length 1677 bp, GFAP promoter), GFAPCE (length 1061bp, CMV enhancer+gfaABC1D promoter) and GFAP104 (length 848bp, EF1α enhancer+gfaABC1D promoter). Recombinant AAV5 and AAV9 Adeno-associated virus were produced by PackGene® Biotech, LLC, purified through iodixanol gradient ultracentrifuge and subsequent concentration. Viral titers were determined by quantitative PCR-based method. The virus was diluted in PBS containing 0.001% F-68. Unless otherwise specified, AAV titre used in this study was mostly in the order of 10^11^ – 10^12^ GC/ml, with the exception of experiments in Fig. 1 to purposely demonstrate high neuronal leakage using high titre of 10^13^ GC/ml.

### Stereotaxic viral injection

The adult mice were anesthetized with isoflurane and then placed into a stereotaxic frame. The hair of the head was shaved off and a midline scalp incision of the brain was operated, then a drill hole (0.6 mm) on the skull was created for the virus injection. 1 μl virus was injected into the cortex (AP +0.6 mm, ML −2 mm, DV −0.7 mm) and striatum (AP +1.0 mm, ML +2.0 mm, DV −3.0 mm) at the speed of 100 nl/min. After viral injection, the needle was kept in the brain for about 8 mins and then slowly withdrawn.

### In vivo two-photon live imaging

A two-photon microscope (Zeiss LM780, Oberkochen, Germany) equipped with a Ti:Sapphire laser source (120 fs width pulses, 90 MHz repetition rate, Coherent Inc., USA) was used for live imaging. The mouse was held by an in vivo imaging platform (Beijing Xing Lin Biotechnology Co., Ltd.) during the long-term repetitive imaging process. The excitation wavelength was set to 920 nm. Imaging was achieved using a 20X water-immersion objective (N.A. 1.0, Carl Zeiss, Oberkochen, Germany). The image size was 106.27 μm × 106.27 μm or 212.55 μm × 212.55 μm. The image depth was about 100-500 μm from the dura (layer II-V of the cortex).

#### Thinned-skull cranial window preparation and viral injection

Brain surgeries were performed on 4-8 weeks old wild-type mice for viral injection. The mice were anesthetized by intraperitoneal injection of 20 ml/kg 1.25% Avertin (Sigma, T48402). Eye ointment (Guangzhou Baiyunshan Pharmaceutical Co., Ltd.) was applied to cover and protect mouse’s eyes. The hair was shaved over most of the scalp. An incision was made in the scalp and the connective tissue was removed to expose the skull. A small amount of cyanoacrylate glue was placed around the edge of a head-holding adaptor to hold the skull to an immobilization device. Then, immerse the skull with a drop of ACSF before using a high-speed microdrill (Microdrill 78001, RWD Life Science) to thin a circular area of skull over the mouse neocortex under a dissecting microscope (Carl Zeiss Stemi 305, Germany). After removing the majority of the spongy bone, continue skull thinning with a microsurgical blade to obtain a very thin and smooth preparation. After finishing the thinned-skull preparation, we performed viral injection with proper stereotaxic instruments (Model 940, KOPF, USA), an ultra microsyringe pump (UMP3 UltraMicropPump, World Precision Instruments, USA) and a microprocessor-based controller (Micro4, World Precision Instruments, USA). The injection volume and flow rate were 1 μl and 0.1 μl/min. After viral injection, the needle was kept in place for about 10 minutes and then slowly withdrawn. Two-photon imaging began 3 days after viral injection. The thinned skull method for two photon imaging is suitable for retroviral infection.

#### Open-skull cranial window preparation

We employed chronic cranial window preparation for long-term repetitive imaging for the mice injected with AAV. After one day of viral injection, we conducted the cranial window preparation in a vertical laminar flow clean bench (SW-CJ-1FD, AIRTECH System Co., Ltd.). The mice were anesthetized with Avertin. The hair was shaved, and the scalp was washed with iodine tincture and ethanol. A stainless head-holding adaptor was used to immobilize the head with the Cyanoacrylate glue. A hole (2-3 mm diameter) on the skull surface was made by a high-speed microdrill (Microdrill 78001, RWD Life Science) and then covered with a coverslip (Warner, CS-3R). Then, the skull and coverslip were glued by the Vetbond Tissue Adhesive (3M, 1469SB). Lastly, the optical window was fixed to the skull with dental acrylic (Changshu Shang Chi Dental Material Co., Ltd.). To avoid inflammatory response, dexamethasone (~2 mg/kg body weight) was administrated through intramuscular injection for 10 days. For better imaging quality, the mice were given 7-10 days to recover before starting two-photon imaging experiments.

### Immunofluorescence

For brain slice staining, the mice were anesthetized with 1.25% avertin and then perfused, first with saline solution and then with 4% paraformaldehyde (PFA). Brains were isolated and post-fixed overnight with 4% PFA and then incubated in 30% sucrose at 4°C until sank. The tissue was sectioned using the cryostat at 30 um thickness and washed three times in the PBS for 10 mins each. After incubated in in the blocking buffering (10% Donkey serum + 0.3% Triton X 100) for 1.5 h at room temperature, the slices were incubated with the primary antibodies at 4 °C for 48 h (GFP, chicken, abcam, 1:2000; GFAP, rat, Invitrogen, 1:1000; NeuN, guinea pig, Millipore, 1:2000; Sox9, rabbit, Millipore, 1:1000; NeuroD1, rabbit, Abcam, 1:500; RFP, rabbit, Abcam, 1:1000). The brain slices were washed three times in the PBS and then incubated with the secondary antibodies conjugated to Alexa Fluor 488, Alexa Fluor 555, or Alexa Fluor 647 (1:1000) supplemented with DAPI for 1.5 h at room temperature, followed by extensive washing with PBS. The samples were finally mounted with antifade medium and sealed with nail polish. Images were captured using a Zeiss confocal microscope LSM700 or Zeiss Axio imager Z2 microscope.

### Statistical analysis

Quantified data were expressed as mean ± S.D. from three to six mice. Statistical analysis was performed by the unpaired Student’s *t*-test or One-way ANOVA analysis with Tukey post hoc test using GraphPad Prism 7.0. Statistical significance was determined at p value < 0.05.

## Supporting information

Figure S1. In vivo two-photon live imaging of NeuroD1-induced astrocyte-to-neuron conversion at 15 dpi, related to figure 6H

Figure S2. In vivo two-photon live imaging of NeuroD1-induced astrocyte-to-neuron conversion at 35 dpi, related to figure 6H .mp4

## Acknowledgement

This study was supported by Yi-Liang Liu Endowment Fund from Jinan University Education Development Foundation. It was also supported by the National Natural Science Foundation of China (U1801681 to G.C., and 32100793 to Z.-Q.X.) and the Guangdong Province Science and Technology Project (2018B030332001 to G.C.). We would like to thank Chen lab members for rigorous discussion throughout this project.

## Author contributions

G.C. conceived and supervised the entire project, analyzed the data, and wrote the manuscript. L.X. performed the major part of the experiments and data analysis. Z-Q.X. contributed the data of two-photon imaging and the lineage conversion in Aldh1l1-CreERT2xAi14 transgenic mice. Y-W.G. performed the experiment of different titre of AAV. Y-G.X. and M-H.L. performed the conversion studies in the striatum. M-H.L. participated in the result analysis of two-photon imaging. W-Y.J. and S.H. participated in animal surgery, immunostaining and some data analysis. Z.W., W.L. and W-L.L. contributed to the experimental design and data analysis, as well as edited the manuscript.

